# RGS10 differentially modulates NFκB subunit transcription and inflammatory cytokine profiles in peritoneal macrophages

**DOI:** 10.64898/2026.02.09.704897

**Authors:** Janna E. Jernigan Posey, Kruthika Dheeravath, Cassandra L. Cole, Noelle K. Neighbarger, Kelly B. Menees, Malú Gámez Tansey

## Abstract

Regulator of G-protein signaling 10 (RGS10) has been shown to regulate multiple inflammatory pathways relevant to disease pathogenesis. Of particular importance is the ability of RGS10 to negatively regulate the NFkB pathway, a prominent pro-inflammatory pathway implicated in multiple inflammatory disease phenotypes. However, the exact mechanism by which RGS10 regulates NFkB is unknown. Considering that RGS10 translocates into the nucleus upon stimulation, we hypothesize that RGS10 may regulate NFKB through transcription. To determine whether RGS10 mediates NFkB transcription, we stimulated RGS10 KO and B6 peritoneal macrophages and collected cell lysate over 24 hours to assess transcript levels of NFkB and related proinflammatory cytokines. Here we found that RGS10 differentially regulates the transcription of NκKB subunits and NFκB-dependent cytokines. Further studies are warranted to understand the potential role of RGS10 in transcriptional regulation of inflammatory states.

## Introduction

Regulator of G-protein signaling 10 (RGS10) has been shown to regulate inflammatory responses that are protective in multiple chronic inflammatory disease models such as rheumatoid arthritis (RA), metabolic dysfunction, obesity, colitis, periodontist, influenza infection, cancers, and even neurodegenerative disease models (1-9). Mechanistically, RGS10 modulates multiple pathways that regulate the production of pro-inflammatory cytokines in myeloid cells including Stromal interaction molecule 2 (STIM2) mediated calcium entry, glycolytic production of reactive oxygen species (ROS), and suppression of Nuclear Factor kappa-light-chain-enhancer of activated B cells (NFkB)(7, 10, 11). Importantly, NFkB, a master transcriptional regulator of pro-inflammatory cytokines, is translocated to the nucleus upon inflammatory stimulation in myeloid cells allowing for the production of key inflammatory mediators such as Tumor necrosis factor (TNF), Interleukin 1-beta (IL-1B), Interleukin 6 (IL-6), Interleukin 12p40 (IL-12p40), and cyclooxegenase-2 (COX2) (12, 13). Dysregulated NFkB signaling has been identified as an important mediator of multiple chronic inflammatory diseases such as RA, Irritable bowel disease, type 1 diabetes, ect.(12). Specifically, RGS10 deficiency enhances NFkB subunits protein levels upon inflammatory stimulus and also amplifies NFKB activity and proinflammatory cytokine secretion(4, 7). However, the mechanism by which RGS10 regulates NFKB levels is still unknown. Immunoprecipitation assays have further revealed that RGS10 does not have a direct protein to protein interaction with NFkB(10). Interestingly, both RGS10 and NFKB translocate to the nucleus upon inflammatory stimulation indicating that RGS10 may play a nuclear role in regulating the level and activity of NFkB(9, 12). Moreover, immunoprecipitation assays revealed that RGS10 does interact with DNA binding proteins indicating that it may play a role in transcription(10).

Therefore, we hypothesized that RGS10 transcriptionally regulates NFKB. To test this, we analyzed NFkB and proinflammatory transcripts of lipopolysaccharide (LPS) stimulated wild type C57B6/J (B6) and RGS10 Knock out (KO) peritoneal macrophages (pMacs) at 1,2,4,8, and 24h. Importantly, animals were given opioid analgesics prior to pMac collection. Under these conditions, we found that RGS10 differentially regulates the transcription of NFKB subunits and NFKB-dependent cytokines. Specifically, we demonstrate that RGS10 does not regulate the kinetics of NFKB subunit transcription or IL-1B cytokine transcription levels. However, we do see that RGS10 deficiency briefly results in higher levels of p65 transcript levels shortly after initial stimulation. Moreover, we found that RGS10 KO pMacs produced significantly less TNF transcript than their B6 counterparts and this difference was driven by male mice. We also found no difference in secreted protein levels of TNF, IL-1β, or IL-10 but did see elevated levels of IL-6, IL-12, and KCGRO in LPS stimulated RGS10 KO pMacs. In summary, we demonstrate transient and differential transcriptional regulation of NFKB pathway components by RGS10. These data warrant further investigation into the potential role of RGS10 in transcriptional regulation. Importantly, these phenotypes we collected with the use of opioid analgesics and therefore our data demonstrate that RGS10 can regulate differential myeloid pro-inflammatory profiles under the influence of such opioid analgesics.

## Materials and Methods

### Animals

Mice were housed in the McKnight Brain Institute vivarium at the University of Florida and maintained on a 12:12 light–dark cycle with *ad libitum* access to water and standard rodent diet chow. All animal procedures were approved by the Institutional Animal Care and Use Committee and followed the Guide for the Care and Use of Laboratory Animals from the National Institutes of Health at the University of Florida. Generation of the RGS10 Knock out (KO) line onto a C57B6/J background has been previously described(7). 4 to 5 month-old male and female C57B6/J and RGS10 KO mice (n=12/group) were given 50uL of sustained-release Buprenorphine (ZooPharm) subcutaneously 30 minutes prior to intraperitoneal injection of 1mL of 3% thioglycolate. 3 days later animals were sacrificed via cervical dislocation for peritoneal macrophage collection.

### Peritoneal macrophages

Peritoneal macrophage(pMacs) collection occurred as previously published(14). Specifically, Pmacs collection occured in a laminar flow hood. After cervical dislocation and spraying down the animals with 70% ethanol, 10mL of cold RMPI were injected into the peritoneum. Peritoneal lavage fluid was collected, passed through 70 uM filters prewet with 5mL of HBSS-/-(Gibco, 14175103) into 50mL conical tubes. Filters were rinsed with 5mLs of HBSS-/-(Gibco, 14175103) twice and centrifuged at 400xg for 5 minutes at 4°C. If blood cells were present in the sample, cells were treated with 1mL of ammonium-chloride-potassium lysis buffer for 1min and then pelleted at 400xg for 5 minutes at 4°C. Cells were taken forward and processed in the biosafety hood using sterile technique. Supernatant was aspirated off cells, and cells resuspended in 3mL of warm plating media (RPMI, 10% fetal bovine serum, and 1% penicillin/streptomycin). 10uL of cells were taken forward for counting using a 1:1 diluted trypan blue (1/4th trypan blue in sterile 1XPBS) stain and the CountessTM. Cells were plated at 1X106 density in a 2mLs of culture media in a 12 well plate. Cells were placed in the incubator at 37° C and 5% CO2 to rest for 2 hours. After 2 hours, cells were washed with sterile DPBS (Gibco, 14190235) to remove non-adherent cells and new pre-warmed culture media (RPMI supplemented with 10% FBS and 1x Pen-Strep) was added. Cells were treated with either vehicle or 100ng/mL LPS for 1,2,4,8, or 24hrs. Media was collected and flash frozen in liquid nitrogen. Cells were lysed with 350uL BME and RLT lysis buffer (20uL of BME per 1mL RTL buffer) and mechanically homogenized by scrapping each well with a cell scraper.

### RNA extraction

Bench areas were cleaned with RNase Zap prior to RNA extraction. Lysed pMac cells were processed for RNA extraction via RNeasy Kit (Qiagen) according to manufacturer’s protocol. Briefly, lysed solution was transferred to Qiashredder tubes for complete homogenization. Qiashredder flow-though was stored in -80°C until sample batches were collected. Samples were thawed and mixed with 350uL of 70% ethanol and transferred to RNAeasy spin columns. Samples were centrifuged for 15 seconds at 10,000rmp(9,391xg) at room temperature. Flow-through was discarded and nucleic acids trapped in the RNAeasy membrane were washed with 700uL of RW1 buffer and 500uL of RPE buffer twice, centrifuging for 15 seconds at 10,000rmp at room temperature and discarding flow-through after each wash. Samples were then centrifuged for 2 minutes at full speed to dry before eluting samples with 30uL of RNAse free water in a 1-minute spin at 10,000rpm. RNA concentrations were determined using Denovix spectrophotometer and samples were stored at -80°C prior to cDNA synthesis.

### cDNA synthesis

DNA was removed from RNA samples prior to cDNA synthesis by diluting RNA samples with nuclease free water and 3.68ul of DNAse master mix (3.36ul of 25mM Mgcl2 + 0.32ul of 1/5 DNAse 1) reaching a total volume of 10ul per sample. Samples were then run through the thermocycler at 37° C for 30 minutes followed by 75° C for 10 minutes and a 4° C hold. 0.4ug of cDNA for each sample was synthesized from RNA using the High-Capacity cDNA Reverse Transcription Kit (Thermo Fisher Scientific, 4368814) according to manufacturer’s instructions. Briefly, 10ul of reverse transcription master mix, which was comprised of 2ul of 10XRT Buffer, 0.8 25XdNT mix (100mMm, 2ul of 10XRT random Primers, 1ul of Rnase Inhibitor, 1ul of Reverse Transriptase, and 3.2ul of Nuclease-Free water per sample, was added to each DNA-free sample. Samples were vortexed and spun down and run in the thermocycler at 25° C for 10 minutes followed by 37° C for 2 hours then 85° C for 5 minutes and a 4° C hold. cDNA samples were then diluted to the final concentration with nuclease-free water and stored at -20 ° C.

### Real Time QPCR

cDNA was combined with SYBR green reagents (Thermo Fisher Scientific, A46112) and specific IDT Forward and Reverse primers at 10ng/well reactions in a 384 well plate in triplicate for Real Time Quantitative PCR analysis in comparative CCT run on the Quant Studio 5 at a volume of 20ul/well. Primer sequences were as follows: mouse NFKB subunit p65 forward: GGA TCC AGT GTG TGA AGA AG and reverse: CTC CTC TAT AGG AAC GTG AAA G, mouse NFKB subunit p105 forward: GGG ACA GTG TCT TAC ACT TAG and reverse: GTC ATC AGA GAT CAA ACC AGA, mouse RGS10 forward: TTG GCT AGC GTG TGA AGA TTT C and reverse: TGG CCT TTT CCT GCA TCT G, mouse TNF forward: CTG AGG TCA ATC TGC CCA AGT AC and reverse: CTT CAC AGA GCA ATG ACT CCA AAG, mouse IL-1B forward: CAA CCA ACA AGT GAT ATT CTC CAT G and reverse: GAT CCA CAC TCT CCA GCT GCA. All QPCR samples were normalized to their own Beta-Actin CT and then to the WT B6 saline group. Samples with an average triplicate variance of ≥ 0.3, had outlying triplicate removed and if samples average remaining variance still exceeded 0.3 the sample was excluded from analysis.

### Mesoscale Discovery Multiplexed Immunoassays

Protein concentrations of TNF, IL-6, IL-1β, KC/GRO, IL-4, IL-10, IFN-Y, IL-5, IL-2, and IL-12p70 in conditioned media were analyzed (MSD, K15048D-2) using multiplexed immunoassays on the MesoScale Discovery (MSD) platform. Samples were plated in duplicate at a 1:1 dilution with MSD diluent 41. Samples were processed according to the manufacturer’s instructions as previously described(14). Plates were analyzed on the MSD Quickplex machine and MSD software (Discovery Workbench Version 4.0). Cytokines with values below the lower limit of detection (LLOD) were not quantified.

### Statistics

Analyses were performed with Graph-Pad Prism 10.6. Group differences were analyzed using Ordinary three-way Anova corrected for multiple comparisons with Tukey post hoc test. Differences between groups across time were analyzed using one-way ordinary ANOVA corrected for multiple comparisons with Tukey’s post hoc analysis. Samples that are statistically different do not share the same letter. p values ≤ 0.05 were considered statistically significant.

## Results

To determine whether the RGS10 mediated NFkB-dependent transcription, we stimulated RGS10 KO and B6 peritoneal macrophages and collected cell lysate over 24 hours to and assessed transcript levels of NFkB and related proinflammatory cytokines. If the hypothesis that RGS10 negatively regulates NFkB transcription is correct, we would expect to see that RGS10 deficiency enhances NFkB-dependent pro-inflammatory gene transcription. Here, we find that RGS10 does not differentially regulate the kinetics of NFKB subunit transcription when analyzed over the course of 24 hours, nor did we find an enhancement in the transcription of NFkB-dependent genes such as TNF and IL-1B (Figure 1). However, we did find that RGS10 deficiency results in higher levels of the p65 NFkB subunit transcript at 2 hours post LPS stimulation (Supplemental figure 1D), indicating a potential time sensitive role for the regulation of the p65 NFkB subunit by RGS10. Importantly, we demonstrate that RGS10 transcript levels are reduced upon stimulation and recover over time, which has been previously reported by multiple studies (Figure 1B)(9, 15). Interestingly, we found a genotype effect of TNF transcripts indicating that RGS10 KO peritoneal macrophages (pMacs) were producing less TNF than their B6 counterparts (Figure 1C).

**Figure 1.**
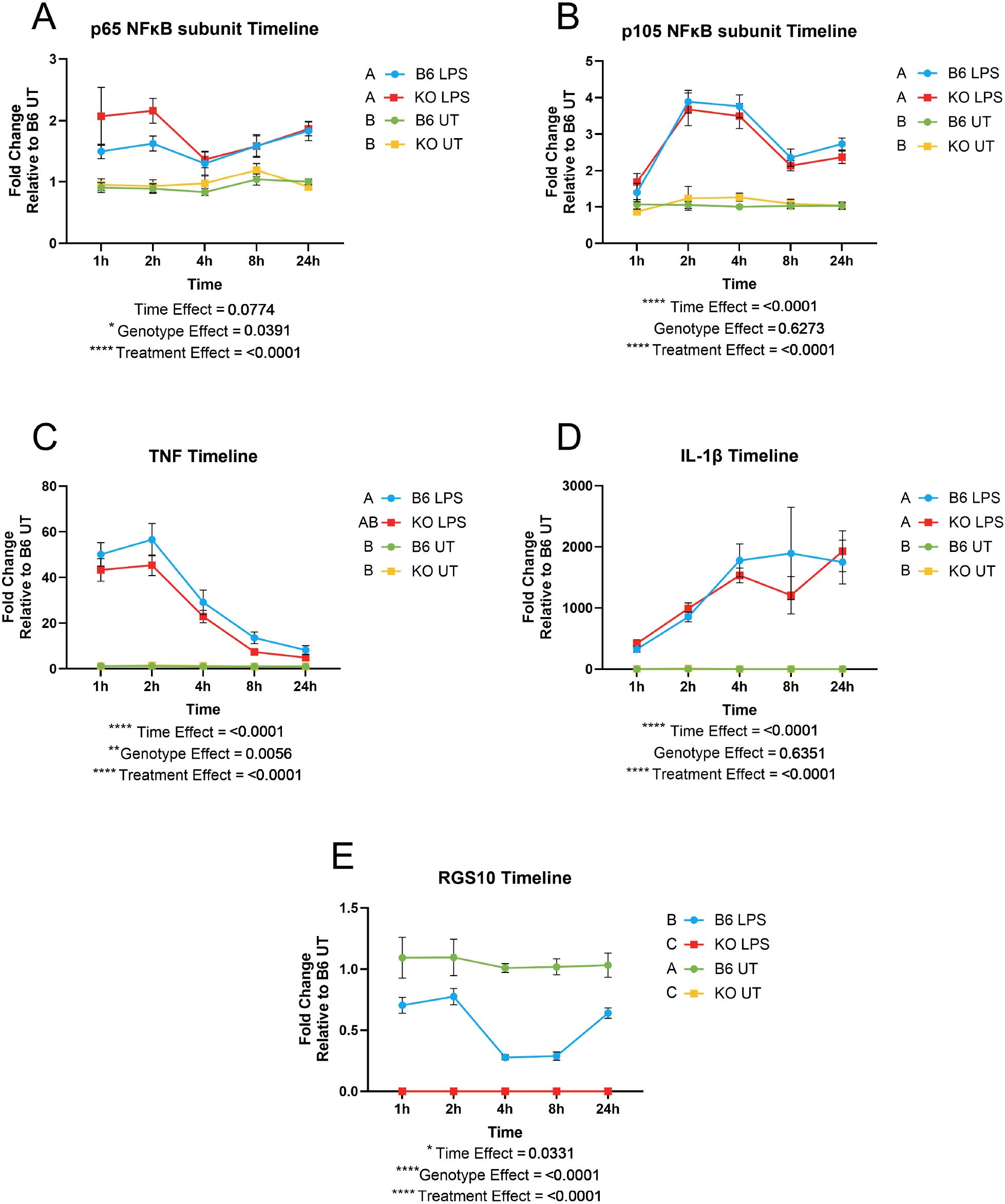
RGS10 differentially regulates the transcription of NFκB subunits and NFκB-dependent transcripts. The red line represents RGS10 KO pMacs treated with LPS, blue line represents B6 pMacs treated with LPS, yellow line represents RGS10 KO pMacs treated with vehicle, and the green line represents B6 pMacs treated with vehicle. A) Fold change of p65 NFκB transcripts relative to control over 24h. B) Fold change of p105 NFκB transcripts relative to control over 24h C) Fold change of TNF transcripts relative to control over 24h. D) Fold change of IL-1βtranscripts relative to control over 24h. E) Fold change of RGS10 transcripts relative to control over 24h Significant genotype present on appropriate graphs. Samples that are statistically different do not share the same letter.

To understand if any of our results were sex specific, we analyzed our data in males and females separately. Here we found that the genotype effect on TNF is only present in male RGS10 KO pMacs and not females (Figure 2C,G). Otherwise, male and female pMacs produced similar transcriptional inflammatory responses to LPS. Interestingly, early transcription of the p65 subunit of NFkB was significantly enhanced with RGS10 deficiency in females but not males (Supplemental Figure 1E-F).

**Figure 2.**
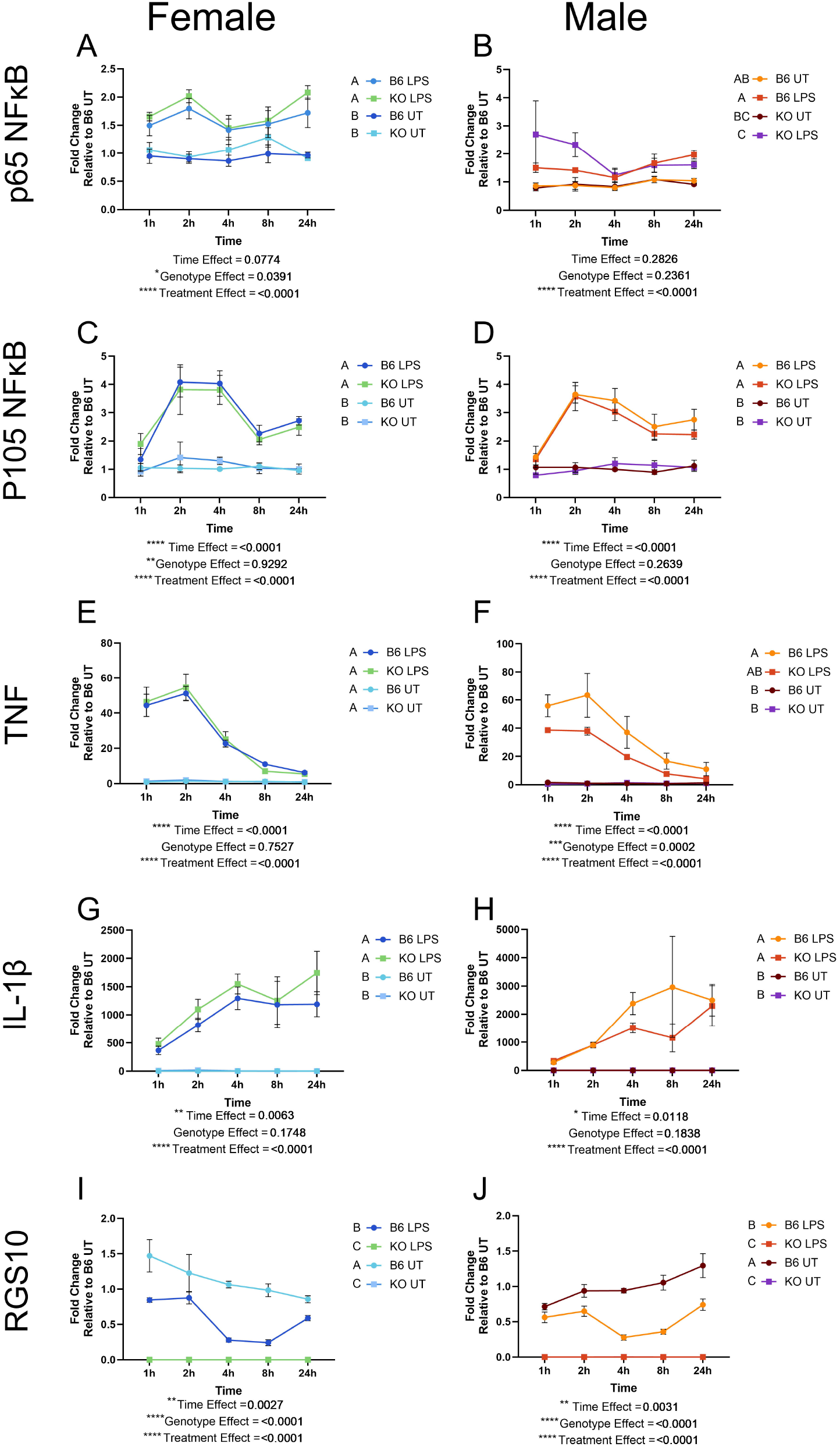
Transcription of the NFκB subunits and pro-inflammatory cytokines is similar across both sexes. Fold change of p65 NFκB transcripts relative to control over 24h in females (A) and males (B). Fold change of p105 NFκB transcripts relative to control over 24h in females (C) and males (D). Fold change of TNF transcripts relative to control over 24h in males (E) and females (f). Fold change of IL-1βtranscripts relative to control over 24h in males (G) and females (H). Fold change of RGS10 transcripts relative to control over 24h in males (I) and females (J). Significant genotype present on appropriate graphs. Samples that are statistically different do not share the same letter.

It was surprising that our data did not demonstrate any enhancement in TNF or IL-1β transcripts levels as previous literature reported exacerbated TNF and IL-1β protein levels with RGS10 KO in pMacs(16). Therefore, we examined the level of secreted pro-inflammatory cytokines from the conditioned media. Again, we found that TNF and IL-1β levels did not change with RGS10 deficiency, nor did levels of IL-10 (Figure 3 A,D,F). Instead, we found that RGS10 deficiency exacerbated the levels of IL-6, KCGRO, and IL-12 (Figure 3 B,C,E).

**Figure 3.**
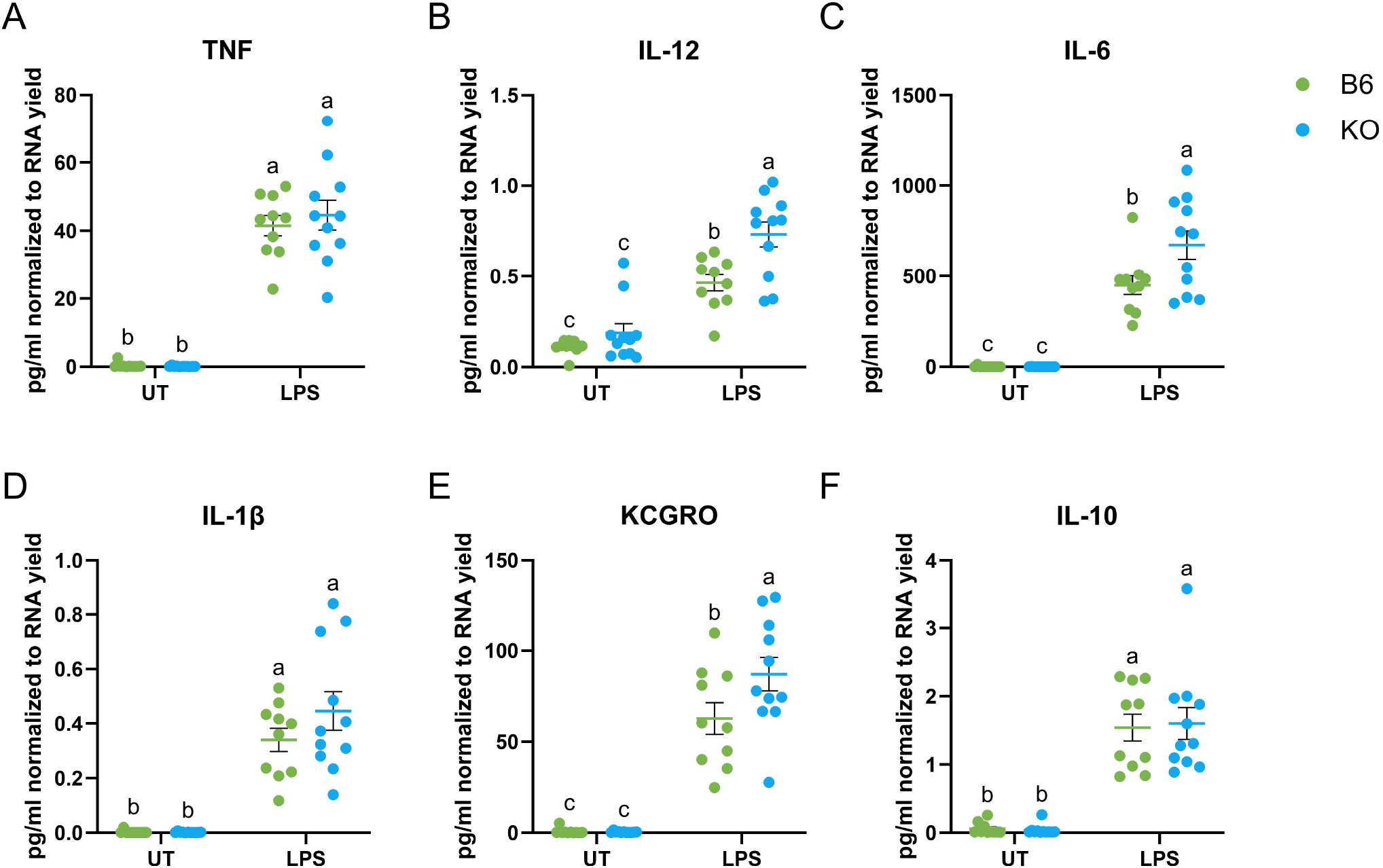
RGS10 regulates the secretion of pro-inflammatory cytokines in peritoneal macrophages Level of pro-inflammatory cytokines from male and female pMacs. Green dots are B6 pMacs and Blue dots are RGS10 KO pMacs. A) pg/mL normalized to RNA yield of TNF protein levels 24h post stimulation. B) pg/mL normalized to RNA yield of IL-12 protein levels 24h post stimulation. C) pg/mL normalized to RNA yield of IL-6 protein levels 24h post stimulation. D) pg/mL normalized to RNA yield of IL-1B protein levels 24h post stimulation. E) pg/mL normalized to RNA yield of KCGRO protein levels 24h post stimulation. F) pg/mL normalized to RNA yield of IL-10 protein levels 24h post stimulation. Samples that are statistically different do not share the same letter.

## Discussion

Overall, our data demonstrates that RGS10 differentially regulates the transcription of NFKB subunits and NFKB-dependent cytokines in peritoneal macrophages. However, this effect is time sensitive and displays multiple sex biases. The brief and not sustained increase of the p65 NFkB subunit transcript levels with RGS10 deficiency can likely be explained by the downregulation of RGS10 transcript post LPS stimulation, rending a lack of RGS10 in which to regulate p65 NFkB transcript levels with past the initial stimulation (∼within 2 hours of stimulation). In support of this idea, RGS10 protein levels have also been observed to decrease with LPS stimulation in other myeloid populations over the course of 24-48 hours prior to returning to baseline levels(9, 15).

Moreover, our data reveals novel cytokine transcription and secretion patterns mediated by RGS10. Specifically, we observe, in contrast to previous studies, that RGS10 deficiency did not impact transaction of key NFκB-dependent cytokine IL-1b and even resulted in the reduction of TNF transcript in a sex-dependent manner. Additionally, we observe, again in contrast to previous studies, no change in TNF, IL-1β, or IL-10 cytokine secretion in RGS10 deficient pMacs compared to their B6 counterparts upon LPS stimulation. Interestingly, IL-6, KCGRO, and IL-12 cytokine secretion was exasperated by RGS10 deficiency instead, capturing a differential inflammatory profile that is regulated by RGS10.Importantly, this differential mediation of inflammation may be due to the use of buprenorphine in our study that previous studies were not stipulated to use(16). Buprenorphine is a partial mu opioid receptor agonist, and the mu opioid receptor, which is expressed in myeloid cells among others, is an inhibitory GPCR(17). RGS10 was canonically identified as GTPase-activating protein with high-affinity for inhibitory GPCRs(18). Therefore, we speculate that the regulation of pro-inflammatory responses by RGS10 may be significantly influenced by the use of opioids. Follow-up studies are necessary to clarify the influence of buprenorphine and other opioids on RGS10-mediated myeloid cell activation. Consequently, our study reveals that RGS10 differentially and transiently regulates the transcription of NFκB pathway components and that RGS10 may mediate alternative inflammatory responses under the influence of opioid analgesics.

## Acknowledgments

The authors would like to thank the Tansey lab for useful discussions in the completion of this paper. For open access, the author has applied a CC BY public copyright license to all Author Accepted Manuscripts arising from this submission.

## Author Contributions

JEJP: Conceptualized and designed the study, completed mouse injections, in vitro work, and biochemical analysis, wrote and edited the original manuscript. KD: Assisted with in vitro work and biochemical analysis. CLC and NKN: Maintained mouse colonies and assisted in tissue harvesting. KBM: Participated in study design, data analysis, and manuscript editing. MGT: Contributed to conception and design of the study as well as preparing the final manuscript.

## Data Availability

The data generated in this study are available at the following Zenodo Repository DOI: 10.5281/zenodo.18319174

## Funding

This work was supported in part by the NINDS T32 Pre-Doctoral Training Program in Movement Disorders and Neurorestoration (5T32NS082168-09), Training Grant on Alzheimer’s Disease and ADRD at Indiana University (5T32AG071444-05) – JEJP and the National Institute of Health and the National Institute of Neurological Disorders and Stroke (Grant RF1NS28800 ) – MGT, and Stark Neuroscience Research Institute (SNRI) at Indiana University School of Medicine (IUSM).

### Abbreviations

BME: Beta mercaptoethanol
cDNA: complementary DNA
Cox2: Cyclooxygenase 2
DNA: Deoxyribonucleic Acid
DPBS: Dulbecco’s Phosphate Buffered Saline
GPCR: G Protein Coupled Receptor
IL-12: Interleukin 12
IL-12p40: Interleukin 12p40
IL-1β: Interleukin 1β
KCGRO: Keratinocyte chemoattractant/Growth-regulated oncogene
KO: Knock Out
LLOD: Lower limit of detection
LPS: Lipopolysaccharide
MSD: Mesoscale Discovery
NFkB: Nuclear Factor kappa-light-chain-enhancer of activated B cells
pMac: Peritoneal macrophages
QPCR: Quantitative real time polymerase chain reaction
RA: Rheumatoid Arthritis
RGS10: Regulator of G Protein Signaling 10
RNA: Ribonucleic Acid
RPMI 1640: Roswell Park Memorial Institute 1640
STIM2: Stromal interaction molecule 2
TNF: Tumor Necrosis Factor
WT: Wild type

