## Supplemental Data for "RGS10 differentially modulates NFκB subunit transcription and inflammatory cytokine profiles in peritoneal macrophages"

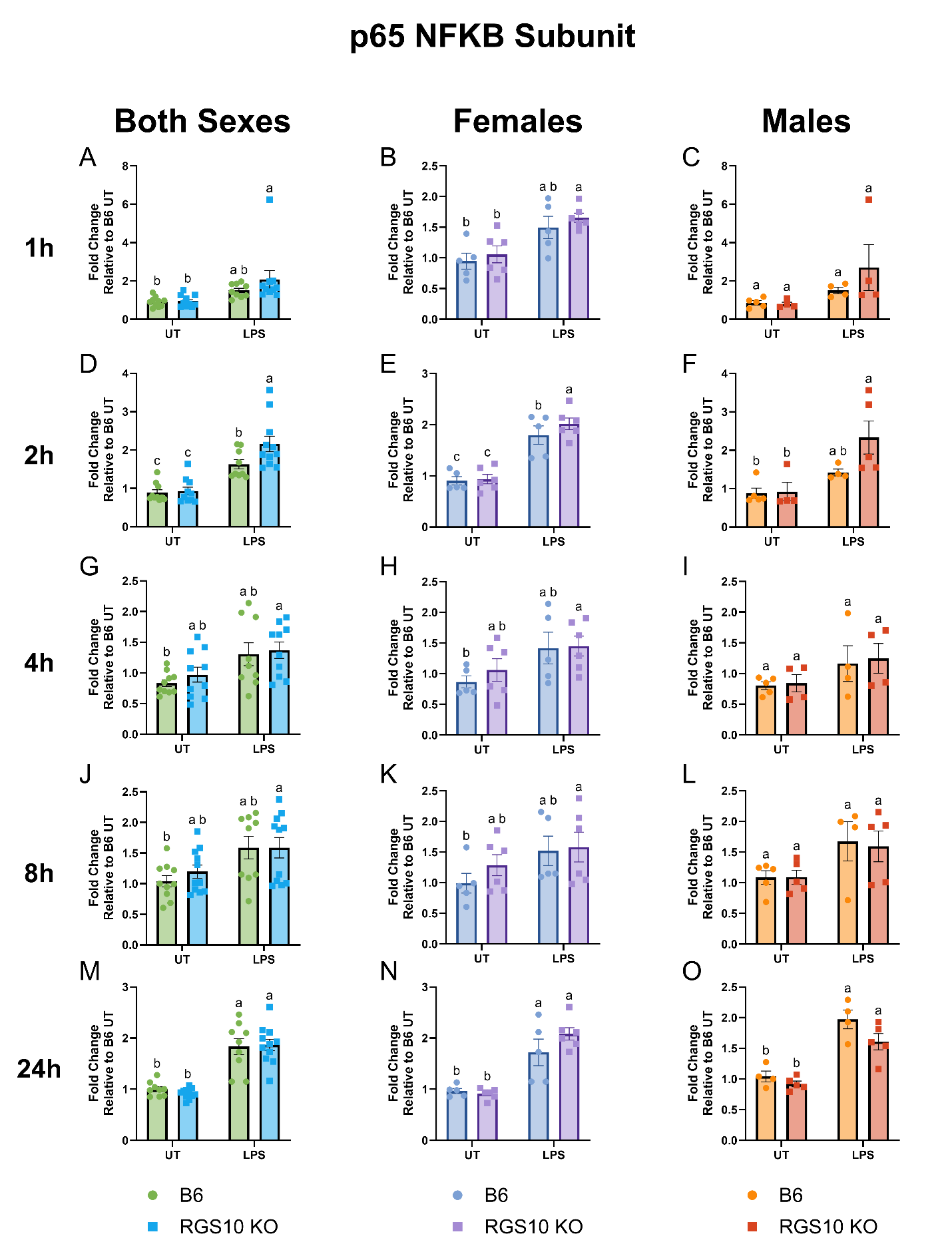

**Supplemental Figure 1: RGS10 suppresses p65 NFKB subunit transcription during initial stimulation response.** Fold change of the transcripts for the p65 subunit of NFKB relative to control at 1h post stimulation in both sexes (A), females (B), and males (C). Fold change of the transcripts for the p65 subunit of NFKB relative to control at 2h post stimulation in both sexes (D), females (E), and males (F). Fold change of the transcripts for the p65 subunit of NFKB relative to control at 4h post stimulation in both sexes (G), females (H), and males (I). Fold change of the transcripts for the p65 subunit of NFKB relative to control at 8h post stimulation in both sexes (J), females (K), and males (L). Fold change of the transcripts for the p65 subunit of NFKB relative to control at 24h post stimulation in both sexes (M), females (N), and males (O). Samples that are statistically different do not share the same letter. Group differences were analyzed using Ordinary two-way Anova corrected for multiple comparisons with Tukey post hoc test. p values ≤ 0.05 were considered statistically significant.

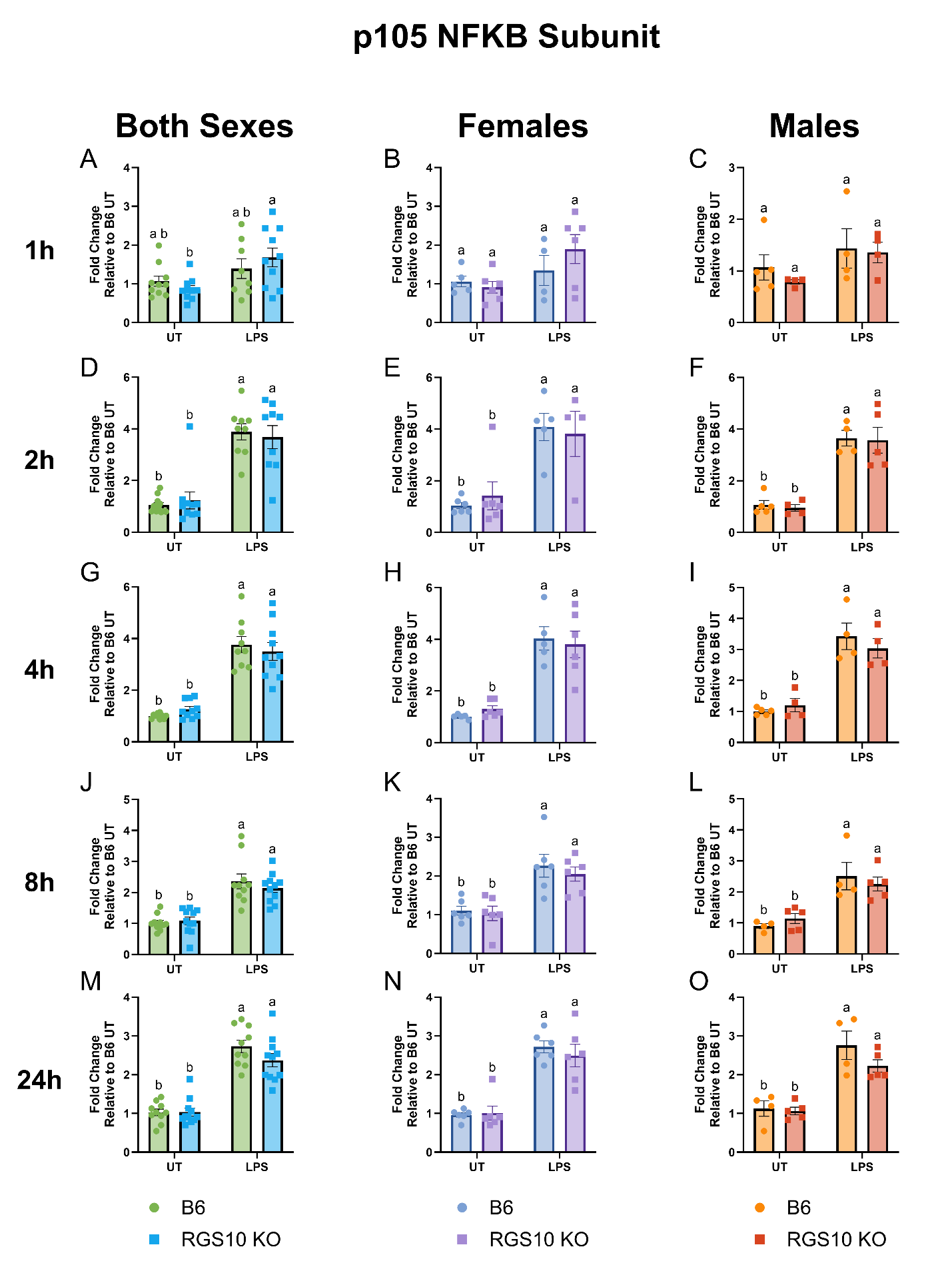

**Supplemental Figure 2: RGS10 dose not modulate p105 NFKB subunit transcription.** Fold change of the transcripts for the p105 subunit of NFKB relative to control at 1h post stimulation in both sexes (A), females (B), and males (C). Fold change of the transcripts for the p105 subunit of NFKB relative to control at 2h post stimulation in both sexes (D), females (E), and males (F). Fold change of the transcripts for the p105 subunit of NFKB relative to control at 4h post stimulation in both sexes (G), females (H), and males (I). Fold change of the transcripts for the p105 subunit of NFKB relative to control at 8h post stimulation in both sexes (J), females (K), and males (L). Fold change of the transcripts for the p105 subunit of NFKB relative to control at 24h post stimulation in both sexes (M), females (N), and males (O). Samples that are statistically different do not share the same letter. Group differences were analyzed using Ordinary two-way Anova corrected for multiple comparisons with Tukey post hoc test. p values ≤ 0.05 were considered statistically significant.

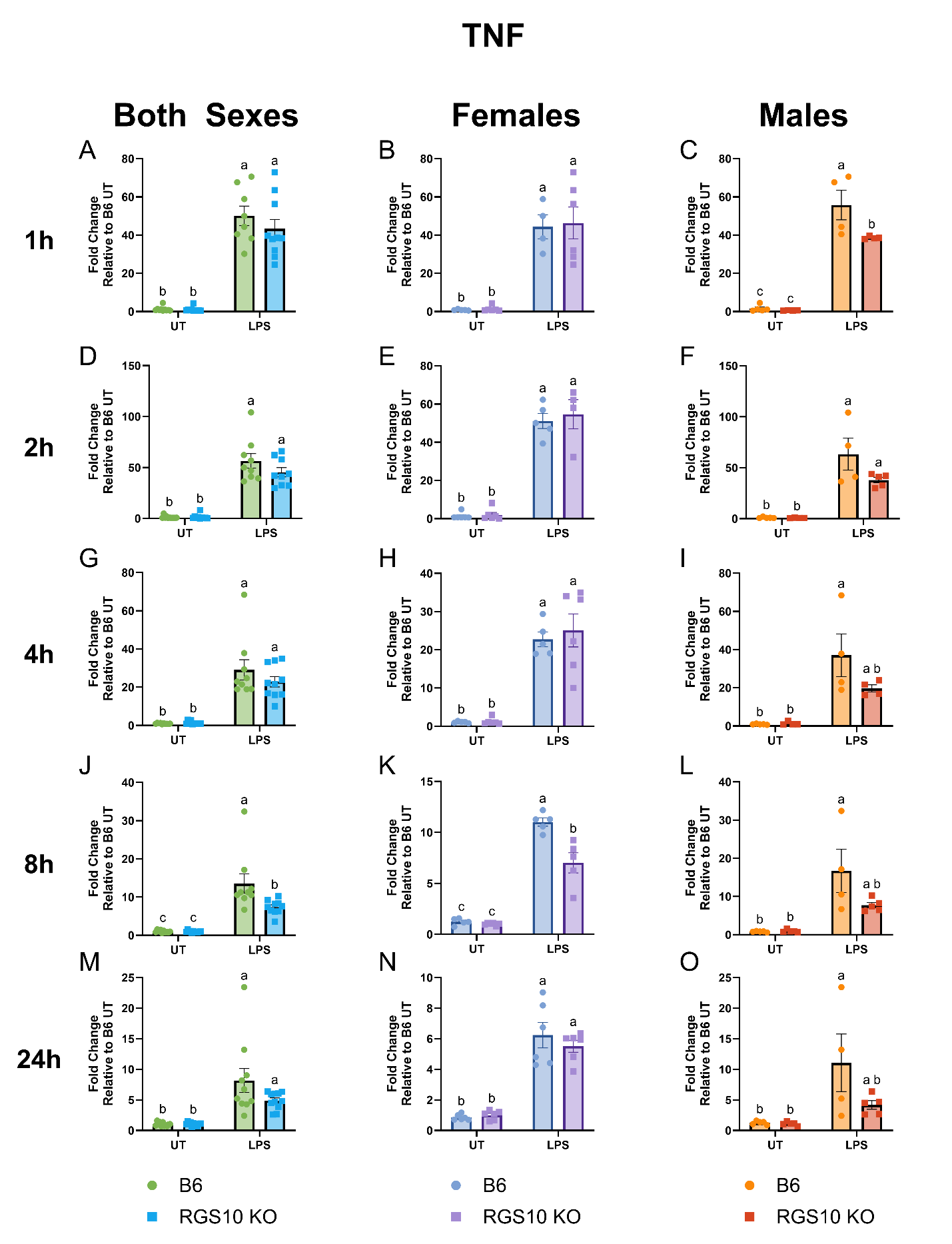

**Supplemental Figure 3: RGS10 deficiency reduces the transcription of TNF over the course of 24h primarily in males.** Fold change of the transcripts for TNF relative to control at 1h post stimulation in both sexes (A), females (B), and males (C). Fold change of the transcripts for TNF relative to control at 2h post stimulation in both sexes (D), females (E), and males (F). Fold change of the transcripts for TNF relative to control at 4h post stimulation in both sexes (G), females (H), and males (I). Fold change of the transcripts for TNF relative to control at 8h post stimulation in both sexes (J), females (K), and males (L). Fold change of the transcripts for TNF relative to control at 24h post stimulation in both sexes (M), females (N), and males (O). Samples that are statistically different do not share the same letter. Group differences were analyzed using Ordinary two-way Anova corrected for multiple comparisons with Tukey post hoc test. p values ≤ 0.05 were considered statistically significant.

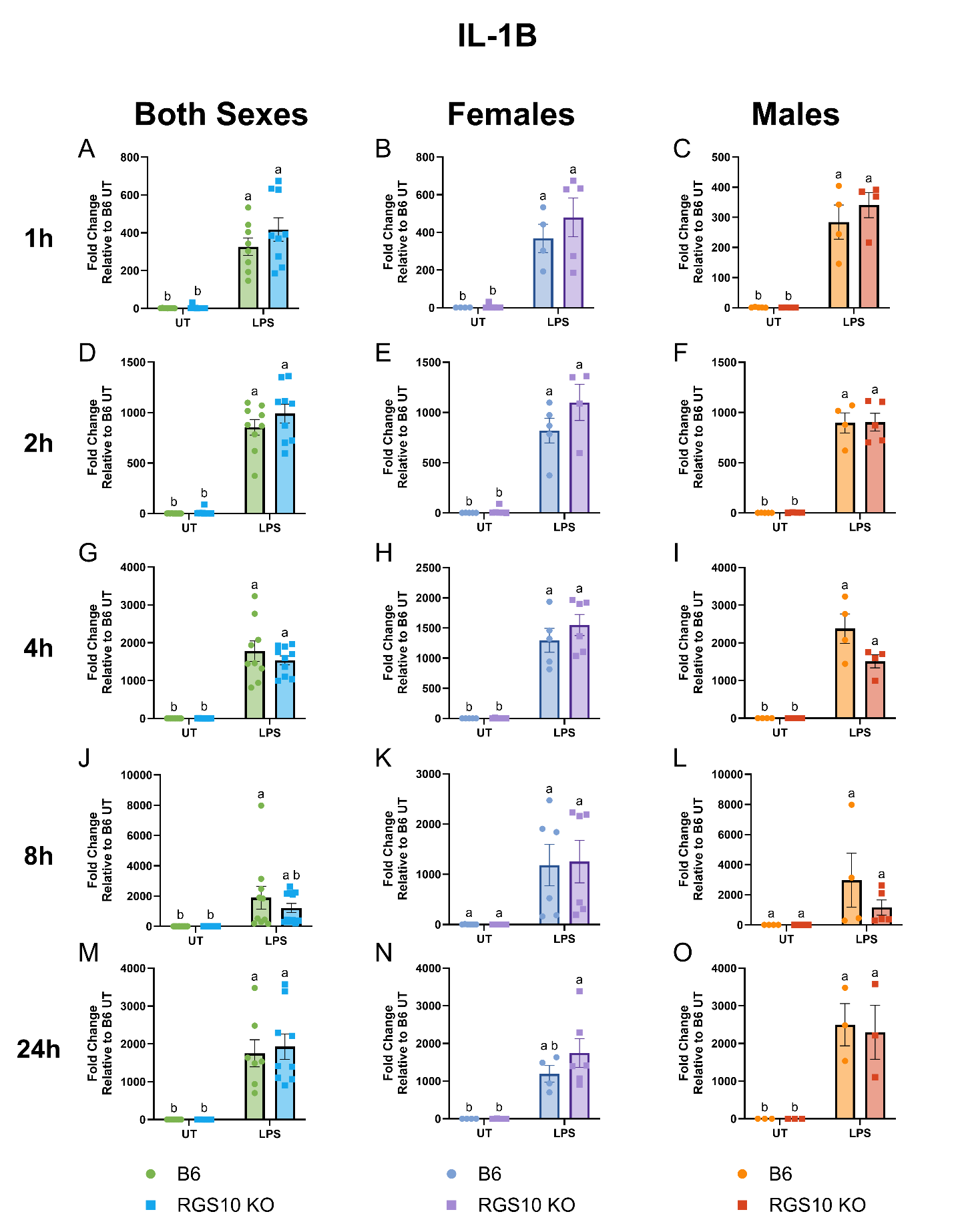

**Supplemental Figure 4: RGS10 does not modulate IL-1B transcription.** Fold change of the transcripts for IL-1B relative to control at 1h post stimulation in both sexes (A), females (B), and males (C). Fold change of the transcripts for IL-1B relative to control at 2h post stimulation in both sexes (D), females (E), and males (F). Fold change of the transcripts for IL-1B relative to control at 4h post stimulation in both sexes (G), females (H), and males (I). Fold change of the transcripts for IL-1B relative to control at 8h post stimulation in both sexes (J), females (K), and males (L). Fold change of the transcripts for IL-1B relative to control at 24h post stimulation in both sexes (M), females (N), and males (O). Samples that are statistically different do not share the same letter. Group differences were analyzed using Ordinary two-way Anova corrected for multiple comparisons with Tukey post hoc test. p values ≤ 0.05 were considered statistically significant.

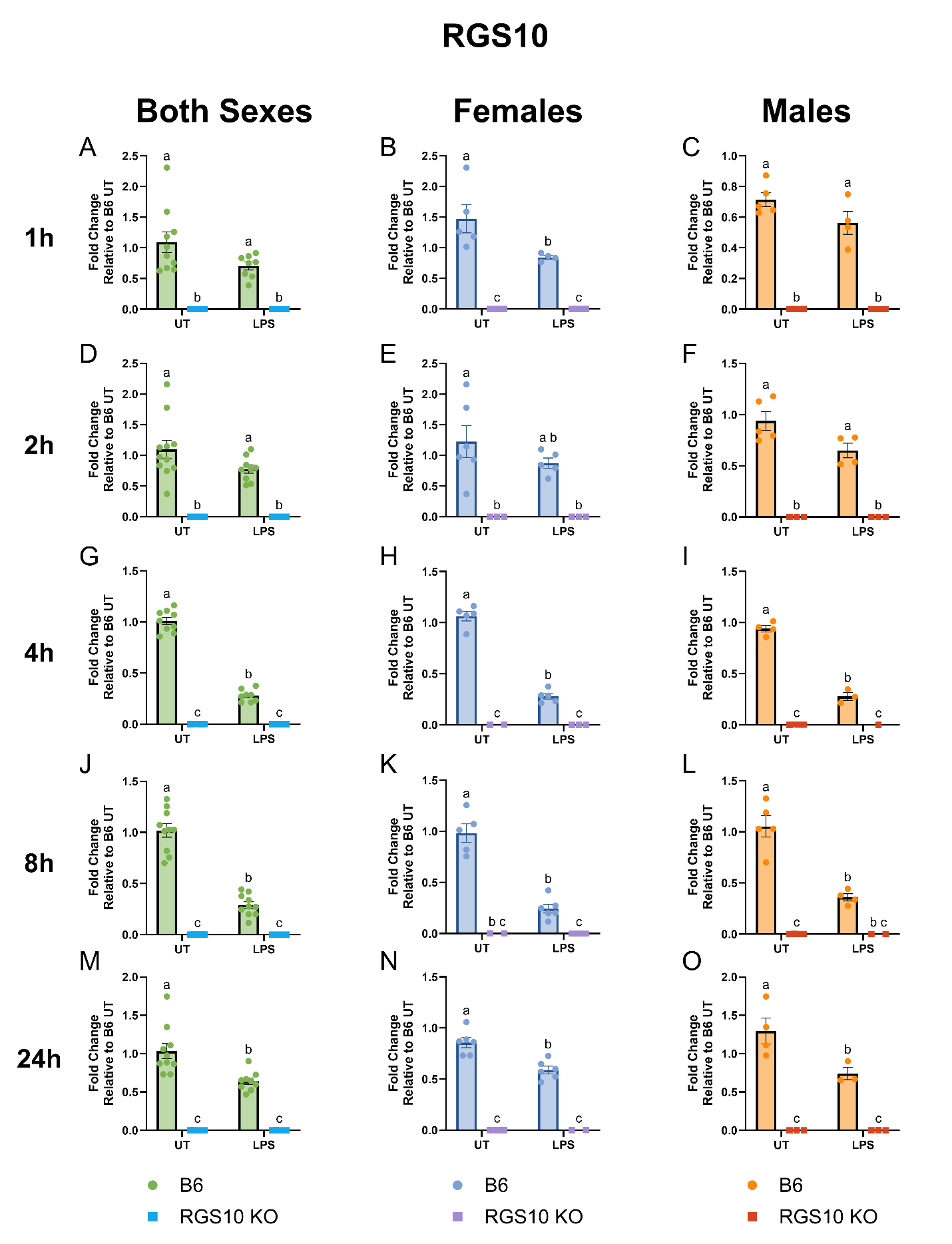

**Supplemental Figure 5: LPS stimulation reduces RGS10 transcription levels over the course of 24h.** Fold change of the transcripts for RGS10 relative to control at 1h post stimulation in both sexes (A), females (B), and males (C). Fold change of the transcripts for RGS10 relative to control at 2h post stimulation in both sexes (D), females (E), and males (F). Fold change of the transcripts for RGS10 relative to control at 4h post stimulation in both sexes (G), females (H), and males (I). Fold change of the transcripts for RGS10 relative to control at 8h post stimulation in both sexes (J), females (K), and males (L). Fold change of the transcripts for RGS10 relative to control at 24h post stimulation in both sexes (M), females (N), and males (O). Samples that are statistically different do not share the same letter. Group differences were analyzed using Ordinary two-way Anova corrected for multiple comparisons with Tukey post hoc test. p values ≤ 0.05 were considered statistically significant.
